# Modulation of decodable semantic features of brain activity via selection attention

**DOI:** 10.1101/2025.06.13.659505

**Authors:** Ayane Okuno, Hiroto Q. Yamaguchi, Shinji Nishimoto, Tomoya Nakai

**Author notes:** Corresponding author, Full postal address: 6F Sanpo Sakuma Building, 1-11 Kandasakumacho, Chiyoda-ku, Tokyo, Japan.

## Abstract

We frequently encounter linguistic information in multiple modalities, such as text and speech, simultaneously. In such cases, we understand the information by selectively attending to one of the modalities. Previous research has shown that selective attention to a specific stimulus modality modulates cortical activity patterns. However, it remains unclear which aspects of linguistic information are selectively modulated by attention and how such modulation influences the decodability of semantic content. To address this question, we constructed decoding models of latent semantic features from the functional magnetic resonance imaging data of six participants in both unimodal (either visual or auditory) and bimodal conditions (simultaneously visual and auditory, with participants attending to a single modality at a time). In unimodal conditions, we successfully decoded the semantic contents from the brain activity. Decodable features were consistent across modalities in both intra-modal and cross-modal decoding. For bimodal conditions, decoding accuracies for the attended stimuli were higher than for the ignored when training and test stimuli belonged to the same modality. Furthermore, decodable features were more consistent across modalities with attended than ignored stimuli in both intra-modal and cross-modal decoding. These results indicate common decodable semantic features regardless of the presentation modality and that selective attention enhances the semantic representations contributing to such decodability.

## Introduction

In everyday life, we encounter substantial linguistic information in multiple modalities, notably vision (text) and audition (speech). Although each can independently convey meaning, selective attention to one modality is often required when receiving simultaneous inputs (Taylor, Lindsay, and Forbes 1967; Massaro and Warner 1977; Spence, Ranson, and Driver 2000). For instance, at a busy train station, an individual may ignore background noise to read the departure time of their train or momentarily shift attention from a smartphone screen to hear the announcement of a delay. These everyday scenarios highlight the critical role of selective attention and modality switching in language processing within complex, multimodal environments.

Previous neuroimaging studies have reported the impact of selective attention on brain representations under multimodal inputs. The brain activity associated with stimuli in the attended modality was higher than that in the ignored modality (Johnson and Zatorre 2006; Mozolic et al. 2008; Keitel et al. 2013; Molloy et al. 2015). Selective attention modulates brain activity on the attended modality also when linguistic information is presented in multiple modalities; such activity is related to the enhanced understanding of attended linguistic information (Mittag et al. 2013; Wang and He 2014; Moisala et al. 2015; Regev et al. 2019; Nakai, Yamaguchi, and Nishimoto 2021). In particular, Nakai, Yamaguchi, and Nishimoto (2021) found that selective attention enhanced the prediction of brain activity in perisylvian regions primarily based on semantic information rather than phonological information. Moreover, the brain activity in these attention-sensitive regions was predictable in a modality-invariant manner, suggesting that modality invariance and attentional selectivity are relevant properties of semantic processing. However, these studies primarily focused on the influence of attention on brain activity. The effects of selective attention on the semantic features associated with brain activity, as well as whether such features are consistent across modalities, remain unclear.

To address these issues, we constructed decoding models able to predict distributed semantic features from brain activity (Pereira et al. 2018; Nishida and Nishimoto 2018; Kivisaari et al. 2019; Fernandino et al. 2022; Frisby et al. 2023) and examined how the decodable semantic features are influenced by presentation modality and selective attention. Distributed semantic features represent words as continuous vectors in a high-dimensional space (Harris 1954; Sahlgren 2008; Boleda 2019), enabling the decomposition of semantic content into fine-grained latent dimensions (Mikolov et al. 2013; Schnabel et al. 2015; Camacho-Collados and Pilehvar 2018). This allowed us to assess which aspects of semantic information are more strongly associated with brain activity under different attentional conditions. We used functional magnetic resonance imaging (fMRI) data from a previous study (Nakai, Yamaguchi, and Nishimoto 2021), where six participants underwent 7-d experiments. The participants were presented with linguistic stimuli either in a single modality or multiple modalities simultaneously. The experiments comprised two parts: (1) in the unimodal experiment, participants were presented with stimuli in a single modality (speech or text). (2) In the bimodal experiment, the participants were simultaneously presented with two stimuli in different modalities (speech and text). They were asked to pay attention to one of the two stimuli and ignored the other.

Decoding models were trained using the data from the unimodal experiment (**Figure 1A**). Semantic features were extracted using Wikipedia2Vec, a neural network model that embeds each word and entity into a vector based on the surrounding words and the page-link information in a Wikipedia corpus (Yamada et al. 2018). We chose this semantic model to compare it with a previous study that constructed encoding models based on the same data (Nakai, Yamaguchi, and Nishimoto 2021). Decoding accuracy was evaluated with left-out data in both unimodal and bimodal conditions (**Figure 1B**). The unimodal analysis served to test whether decodable semantic features are shared across modalities. The bimodal analysis tested how selective attention influences such decodable features. In addition, we evaluated the consistency of both analyses and visualized the cortical voxels contributing to semantic feature decoding.

**Figure 1.**
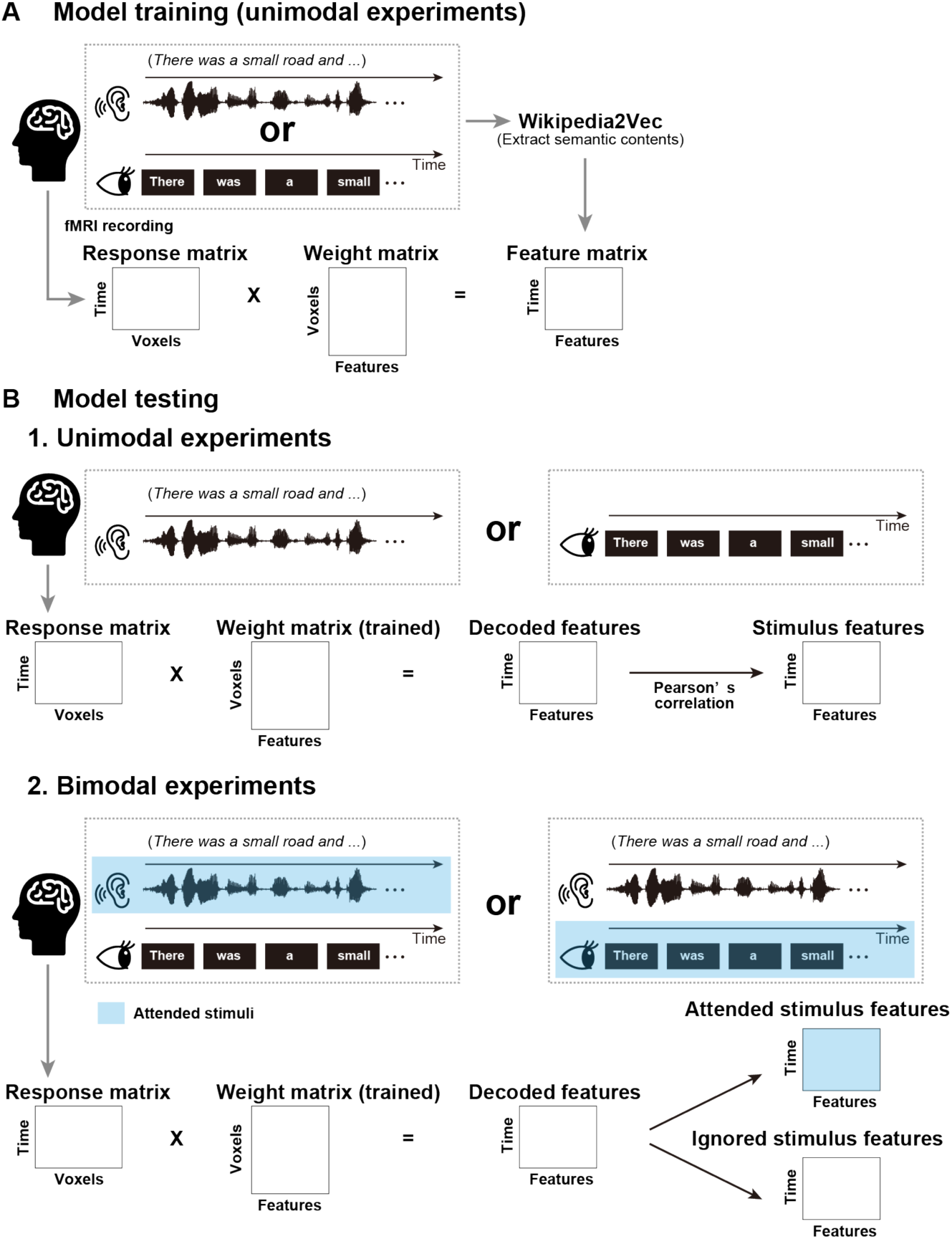
Overview of model training and experimental design. (**A**) For each participant, two distinct models were trained to decode latent semantic features extracted from Japanese speech or texts (referred to as Speech-only and Text-only conditions, respectively) using brain activity. The stimuli shown here are translated in English. Ridge regression was performed to predict semantic features from the brain activity. (**B**) For unimodal analysis, independent test data under the Speech-only or Text-only condition were used. The model output was compared with the semantic features extracted from the actual stimuli using Pearson’s correlation. For bimodal analysis, semantic features were decoded using additional test data, where the participants were simultaneously exposed to speech and text and attended to either one of these modalities (Attend-audio and Attend-visual conditions, respectively). The model output was then compared to the semantic features extracted from attended or ignored stimuli.

## Results

### Semantic features can be decoded in a modality-independent manner

We first tested whether our models could accurately decode semantic features from brain activity in the unimodal experiment. For each participant, we independently trained two decoding models based on one of the two modalities (visual and auditory) and decoded 300 semantic features extracted from the linguistic stimuli of the test data. Depending on the combination of training and test modalities, we obtained four sets of decoding accuracies; two intra-modal conditions (same training and test modalities; auditory-to-auditory and visual to visual) and two cross-modal conditions (different training modality from test modality; auditory to visual and visual to auditory). Over 50% of the features were significantly decoded in all but one condition for one participant (*p* < 0.05, false discovery rate (FDR) corrected; auditory-to-auditory, *r* = 0.125 ± 0.018 (mean ± SD), 78.7% ± 8.9% of features were significant; visual to auditory, *r* = 0.084 ± 0.006, 56.8% ± 6.0% of features were significant; visual to visual, *r* = 0.171 ± 0.042, 85.5% ± 7.8% of features were significant; visual to auditory, *r* = 0.110 ± 0.021, 69.7% ± 9.6% of features were significant; **Figure 2** and **Table S1**).

**Figure 2.**
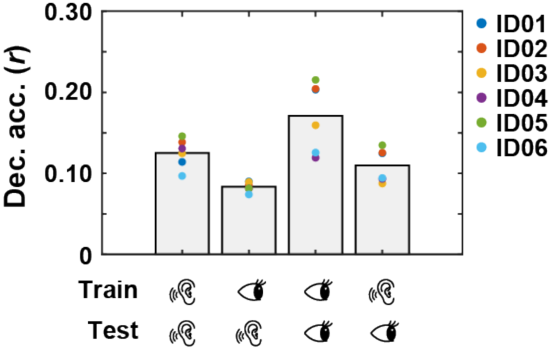
Average decoding accuracies for unimodal conditions. Each data point represents the within-participant average, and the bar plots the between-participant average. Each condition is denoted by icons representing the combination of model training and test modality. First and third bars represent intra-modal conditions, whereas the second and fourth represent cross-modal conditions.

By comparing intra-modal and cross-modal conditions, we examined how the combination of training and test modalities impacts the decoding accuracy. We found that intra-modal conditions exhibited higher decoding accuracy than cross-modal conditions for all six participants for both auditory (Cohen’s *d* = 0.729 ± 0.234) and visual modalities (*d* = 1.034 ± 0.371; **Table S1**). Additionally, we tested whether intra-modal decoding accuracy is affected by the presentation modality. We found a higher decoding accuracy in the visual modality in five out of six participants (Cohen’s *d* = 0.662 ± 0.530; **Table S1**). These results confirm that our models can accurately decode linguistic information from brain activity in both visual and auditory modalities, yet the model performance also depends on the presentation modality.

### Common decodable semantic features across modalities

Next, we examined whether the decodable semantic features are common across modalities. To this end, for each participant, we compared the decoding accuracy of each of the 300 semantic features across conditions, considering all possible combinations across the four conditions. In particular, we separately analyzed four types of combinations: (1) intra-modal conditions, (2) intra-modal and cross-modal conditions with the same test modality, (3) intra-modal and cross-modal conditions with the same training modality, and (4) cross-modal conditions.

First, we compared the decoding accuracies for intra-modal conditions in the visual and auditory modalities (i.e., visual-to-visual and auditory-to-auditory conditions). We found positive correlations between the decoding accuracies in the two intra-modal conditions for all participants (Spearman’s correlation coefficients, *ρ* = 0.642 ± 0.078; **Figure 3A** and **Table S2**). Second, to examine the effect of the training modality, we compared the decoding accuracies between intra-modal and cross-modal conditions with the same test modality but different training modality. Similarly, we found positive correlations for all participants between the decoding accuracies in the visual-to-visual condition and those in the auditory-to-visual condition (*ρ* = 0.733 ± 0.066; **Figure 3B** and **Table S2**), as well as between auditory-to-auditory and visual-to-auditory conditions (*ρ* = 0.595 ± 0.083; **Figure 3C** and **Table S2**). Third, to examine the effect of the test modality, we compared decoding accuracies between intra-modal and cross-modal conditions with the same training modality but different test modality. Again, we found positive correlations for all participants between the decoding accuracies in the visual-to-visual condition and those in the visual-to-auditory condition (*ρ* = 0.439 ± 0.110; **Figure 3D** and **Table S2**), as well as in auditory-to-auditory and auditory-to-visual conditions (*ρ* = 0.573 ± 0.070; **Figure 3E** and **Table S2**). Finally, we compared the decoding accuracies between cross-modal conditions and found positive correlations for all participants (*ρ* = 0.448 ± 0.069; **Figure 3F** and **Table S2**). These results indicate common decodable semantic features across modalities.

**Figure 3.**
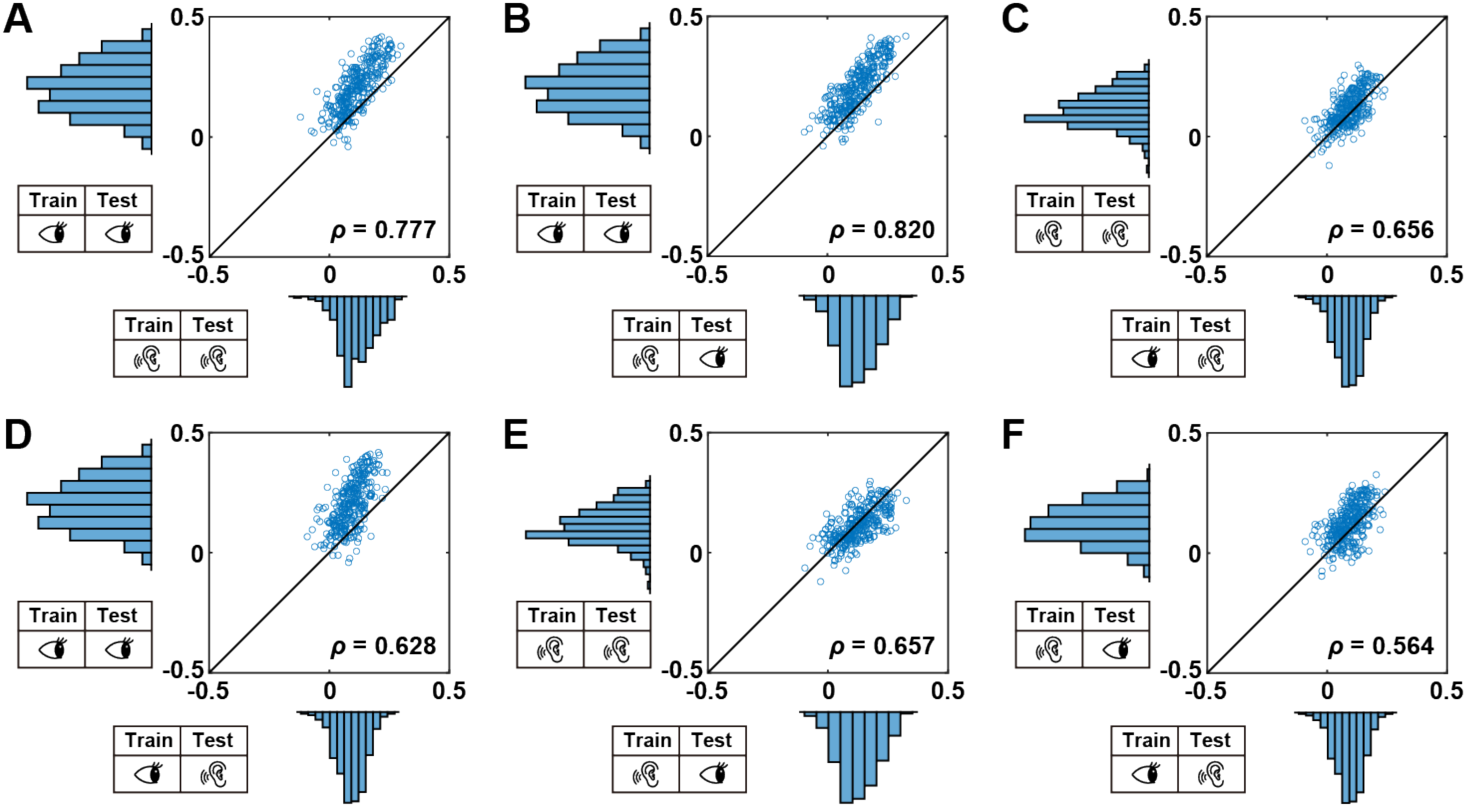
Comparison of decoding accuracies across conditions in unimodal experiments. (**A–F**) Decoding accuracies across conditions for a representative participant (ID05), where each point corresponds to a single semantic feature. (**A**) Comparison among intra-modal conditions in visual and auditory modalities. (**B**) Comparison between intra-modal (visual) and cross-modal conditions (auditory trained and visual tested). (**C**) Comparison between intra-modal (auditory) and cross-modal conditions (visual trained and auditory tested). (**D**) Comparison between intra-modal (visual) and cross-modal conditions (visual trained and auditory tested). (**E**) Comparison between intra-modal (auditory) and cross-modal conditions (auditory trained and visual tested). (**F**) Comparison between the two cross-modal conditions (visual trained and auditory tested vs. auditory trained and visual tested).

### Selective attention increased the decoding performance of semantic features

To investigate how participants’ attention modulates decodable semantic features, we applied our decoding models to the brain activity measured in the bimodal experiment, where participants were simultaneously presented with stimuli in two modalities and paid selective attention (**Figure 4** and **Table S3)**. To minimize the confounding effect of the different modalities, we used the decoding models trained under the unimodal experiment. Over 50% of the features were significantly decoded in all participants in intra-modal conditions with selective attention (*p* < 0.05, FDR corrected; auditory-to-auditory (attend), *r* = 0.138 ± 0.026, 76.7% ± 10.1% of features were significant; visual to visual (attend), *r* = 0.158 ± 0.045, 84.5% ± 7.6% of features were significant). In contrast, most features were not significantly decoded in cross-modal conditions without selective attention (auditory to visual (ignored), *r* = 0.009 ± 0.011, 1.9% ± 1.9% of features were significant; visual to auditory (ignored), *r* = 0.041 ± 0.013, 17.4% ± 15.3% of features were significant). We observed intermediate levels of decoding accuracies in intra-modal conditions without selective attention (visual to visual (ignored), *r* = 0.069 ± 0.022, 45.1% ± 16.3% of features were significant; auditory-to-auditory (ignored), *r* = 0.098 ± 0.016, 62.2% ± 6.4% of features were significant) and cross-modal conditions with selective attention (visual to auditory (attend), *r* = 0.091 ± 0.014, 62.9% ± 9.9% of features were significant; auditory to visual (attend), *r* = 0.065 ± 0.019, 45.7% ± 21.0% of features were significant).

**Figure 4.**
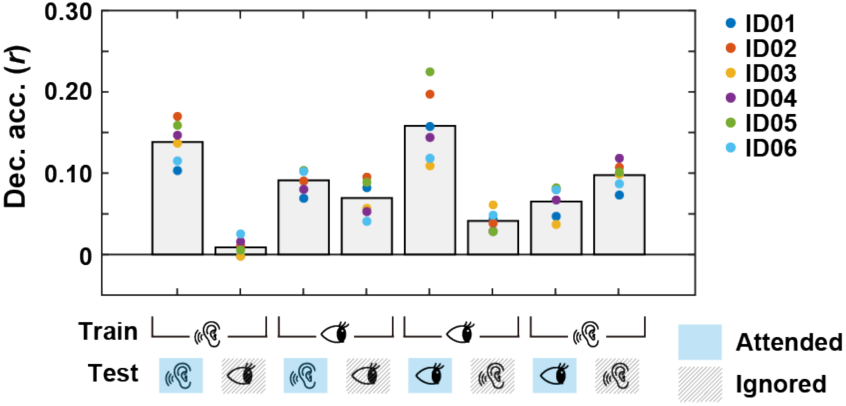
Average decoding accuracies for bimodal conditions. Each data point represents the within-participant average, and bar plots represent the between-participant average. Each condition is denoted by icons representing the combination of model training and test modalities. Stimuli in the attended modality are highlighted in blue.

To replicate the findings in the unimodal data analysis, we directly compared the decoding accuracies between intra-modal and cross-modal conditions. We separately analyzed the decoding results using attended and ignored test stimuli. First, we examined the decoding results of the attended test stimuli. When those stimuli were in the same modality, we found higher decoding accuracy in the intra-modal than in the cross-modal condition for auditory (Cohen’s *d* = 0.830 ± 0.353, 1^st^ and 3^rd^ bar graphs in **Figure 4**) and visually trained models (Cohen’s *d* = 1.335 ± 0.312, 5^th^ and 7^th^ bar graphs in **Figure 4**). We then investigated the decoding results for ignored stimuli. We observed that, similar to attentional stimuli, the intra-modal condition exhibited higher decoding accuracy than the cross-modal condition using auditory (Cohen’s *d* = 0.921 ± 0.351, 2^nd^ and 4^th^ bar graphs in **Figure 4**) and visually trained models (Cohen’s *d* = 0.884 ± 0.279, 6^th^ and 8^th^ bar graphs in **Figure 4**). These results are consistent with the unimodal data analysis, where decoding accuracies were higher in intra-modal than in cross-modal conditions. Meanwhile, the enhanced decoding accuracy in the intra-modal conditions, even for ignored stimuli, suggests that both selective attention and the commonality of the training and test modalities contribute to the decoding accuracy.

We then tested whether selective attention enhances the decoding performance. When the trained modality was the same as the attended modality, decoding accuracies were higher when using features extracted from the attended modality than from the ignored one for all participants for both auditory (Cohen’s *d* = 1.448 ± 0.282; 1^st^ and 2^nd^ bar graphs in **Figure 4**) and visual modalities (Cohen’s *d* = 1.372 ± 0.459; 5^th^ and 6^th^ bar graphs in **Figure 4**). This effect was found regardless of the attended modality, indicating that selective attention promotes semantic processing reflected in brain activity. When the trained and attended modalities were different, we did not find an increase in decoding accuracies by selective attention for the visually trained (Cohen’s *d* = 0.319 ± 0.428; 3^rd^ and 4^th^ bar graphs in **Figure 4**) and auditory trained models (Cohen’s *d* = −0.443 ± 0.264; 7^th^ and 8^th^ bar graphs in **Figure 4**). Accordingly, in the following analyses, we focused on decoding accuracies using the same trained and attended modality (i.e., 1^st^, 2^nd^, 5^th^ and 6^th^ bar graphs in **Figure 4**), as these four conditions are likely to reflect the attentional effect on the decoding performance of semantic information.

### Decodable semantic features are affected by selective attention

To further investigate the effect of attention on decoding accuracy, for each condition, we examined whether selective attention modulates decodable semantic features. In intra-modal conditions, the decoding accuracy of the attended stimuli in the visual and auditory modalities positively correlated for all participants (*ρ* = 0.662 ± 0.088; **Figure 5A** and **Table S4**). When comparing the decoding accuracy of ignored stimuli, a lower correlation was observed for all participants (*ρ* = 0.561 ± 0.118; **Figure 5B** and **Table S4**). In cross-modal conditions, a positive correlation between the decoding accuracy of attended stimuli across modalities was observed for all participants (*ρ* = 0.380 ± 0.060; **Figure 5C** and **Table S4**). Similarly to the previous comparison, when comparing the decoding accuracy of ignored stimuli, the correlation was smaller than that observed in the case of attended stimuli for all participants (*ρ* = 0.104 ± 0.130; **Figure 5D** and **Table S4**). These results indicate common decodable semantic features contributing to the decoding of attended stimuli across modalities but not to that of ignored stimuli.

**Figure 5.**
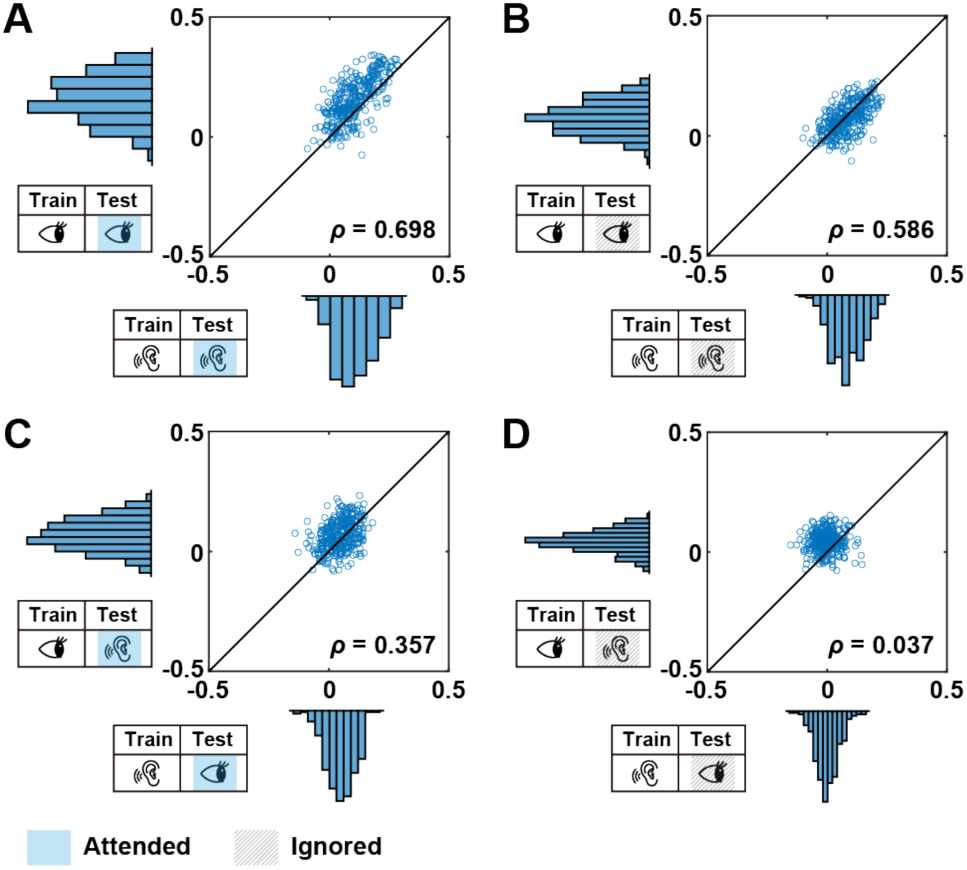
Comparison of decoding accuracies across conditions in bimodal experiments. (**A– D**) Decoding accuracies across conditions for a representative participant (ID05), where each point corresponds to a single semantic feature. (**A**) Comparison of decoding accuracies of attended and (**B**) ignored stimuli in the intra-modal condition. (**C**) Comparison of decoding accuracies of attended and (**D**) ignored stimuli in cross-modal conditions.

### Correspondence between unimodal and bimodal analyses

Unimodal and bimodal experiments provided complementary findings regarding the influence of the training and test modalities and of selective attention, respectively. Additionally, the bimodal experiment demonstrated an interaction effect where decodable features aligned consistently across modalities only when selective attention was focused on linguistic content. Conversely, in the absence of attention, the alignment across modalities decreased.

To further assess the robustness of our findings, we compared the outcomes of unimodal and bimodal data analyses for each participant, considering all possible combinations of training-to-test modalities (i.e., intra-modal and cross-modal) and selective attention (attended and ignored stimuli). Thus, we separately analyzed four types of combinations: (1) intra-modal conditions with attended stimuli, (2) intra-modal conditions with ignored stimuli, (3) cross-modal conditions with attended stimuli, and (4) cross-modal conditions with ignored stimuli.

First, we focused on intra-modal conditions with attended stimuli. The decoding accuracies exhibited positive correlations across all participants in both modalities (auditory modality, *ρ* = 0.619 ± 0.102; **Figure 6A**; visual modality, *ρ* = 0.729 ± 0.092; **Figure 6B** and **Table S5**). Second, we compared the results of the unimodal and bimodal experiments with ignored stimuli, both in the intra-modal conditions. Similarly, the decoding accuracies exhibited positive correlations across all participants for both modalities (auditory modality, *ρ* = 0.585 ± 0.115; **Figure 6C**; visual modality, *ρ* = 0.552 ± 0.140; **Figure 6D** and **Table S5**). Third, we compared the results of the unimodal and bimodal experiments with attended stimuli, in cross-modal conditions. The decoding accuracies exhibited lower correlations in all participants using auditory (visual stimuli; *ρ* = 0.400 ± 0.149; **Figure 6E** and **Table S5**) and visually trained models (auditory stimuli; *ρ* = 0.361 ± 0.069; **Figure 6F** and **Table S5**). Finally, we compared the results of the unimodal experiment with those in case of the ignored stimuli in the bimodal experiment, both in the cross-modal conditions. The decoding accuracies exhibited small correlations using auditory (visual stimuli; *ρ* = 0.092 ± 0.115; **Figure 6G** and **Table S5**) and visually trained models (auditory stimuli; *ρ* = 0.235 ± 0.026; **Figure 6H** and **Table S5**).

**Figure 6.**
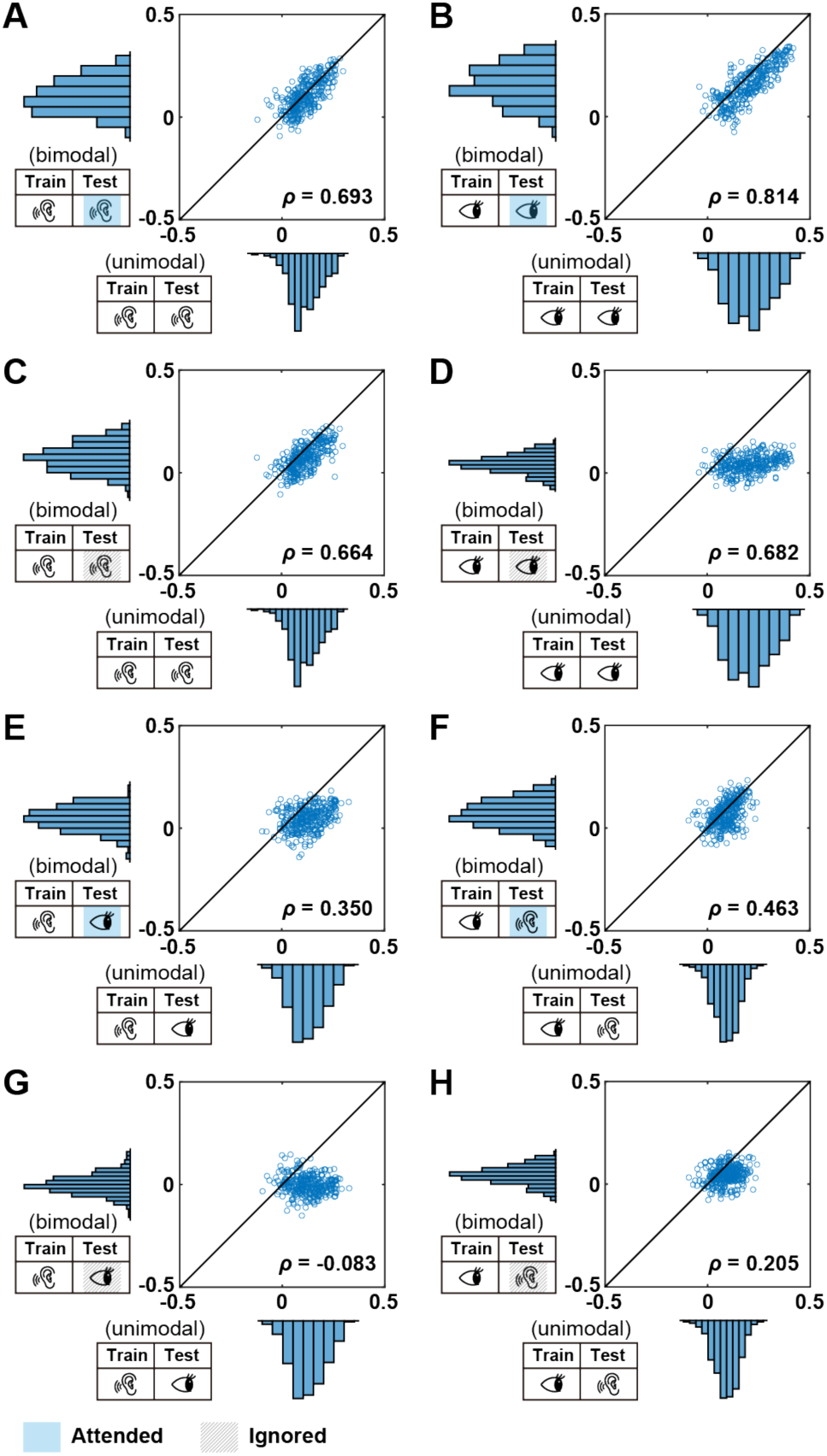
Comparison of decoding accuracy across conditions in unimodal and bimodal experiments. (**A–H**) Decoding accuracies across conditions for a representative participant (ID05), where each point corresponds to a single semantic feature. (**A**) Comparison between unimodal and (attended) bimodal experiments under the auditory intra-modal condition. (**B**) Comparison between unimodal and (attended) bimodal experiments under the visual intra-modal condition. (**C**) Comparison between unimodal and (ignored) bimodal experiments under the auditory intra-modal condition. (**D**) Comparison between unimodal and (ignored) bimodal experiments under the visual intra-modal condition. (**E**) Comparison between unimodal and (attended) bimodal experiments under the cross-modal condition (auditorily trained and visually tested). (**F**) Comparison between unimodal and (attended) bimodal experiments under the cross-modal condition (visually trained and auditorily tested). (**G**) Comparison between unimodal and (ignored) bimodal experiments under the cross-modal condition (auditorily trained and visually tested). (**H**) Comparison between unimodal and (ignored) bimodal experiment under the cross-modal condition (visually trained and auditorily tested).

In sum, under intra-modal conditions, decoding accuracies in unimodal and bimodal experiments were highly correlated regardless of the presence or absence of selective attention. In the cross-modal condition, however, decoding accuracies in unimodal and bimodal experiments more strongly correlated with selective attention to the stimuli. These results suggest consistent decodable semantic features for attended stimuli across modalities, which diversify across modalities without selective attention.

### Brain regions contributing to decoding

To identify brain regions that provide linguistic information relevant to decoding, we analyzed decoder weights and mapped informative voxels onto each participant’s cortical surface. Note that our decoders were trained in unimodal experimental data (i.e., visual and auditory models). We found that the bilateral occipital voxels predominantly contributed to decoding in the visual modality while the bilateral superior temporal lobe contributed to decoding in both modalities surface (**Figures 7A** and **S1**).

**Figure 7.**
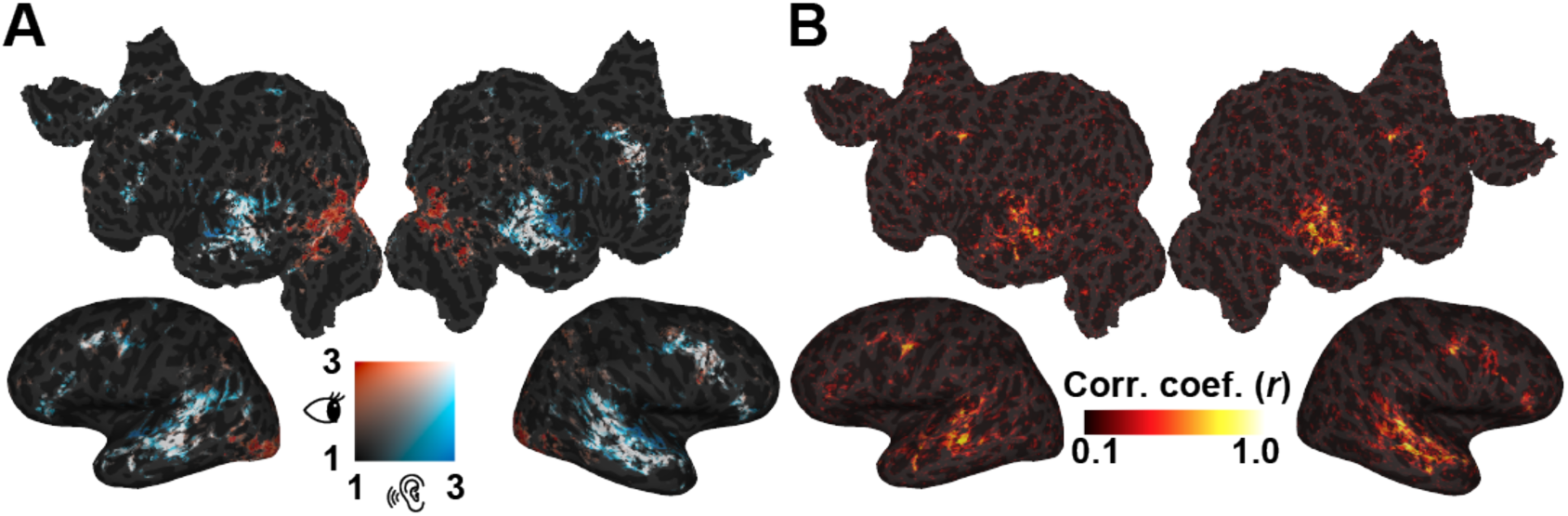
Brain regions contributing to the decoding of semantic features. (A) For visual and auditory models, the maximum absolute values of the model weights are projected onto the cortical surface of a representative participant (ID05). Red voxels represent the visual model and blue voxels the auditory model. (C) Correlation between visual and auditory model weights.

Furthermore, we examined brain regions where similar semantic information is processed across modalities. To this end, we calculated the correlation of weights for each voxel between the visual and auditory models. We found consistently high correlations across participants, particularly in the region surrounding the Sylvian fissure (**Figures 7B** and **S2**). These results are consistent with our previous study on brain regions with shared modality-invariant semantic information (Nakai, Yamaguchi, and Nishimoto 2021).

## Discussion

The current study aimed to examine whether selective attention modulates the semantic features decodable from brain activity and whether such features are shared across modalities. Using semantic decoding models based on unimodal and bimodal experiments, we observed the following results. First, the decoding accuracy was generally higher when training and test stimuli were in the same modality. Second, among 300-dimensional semantic features, there were common decodable features across modalities. Third, for both modalities, the decoding accuracy was higher when selective attention was paid to the target semantic content. Fourth, decodable features were common across modalities under selective attention, but those contributing to the decoding of ignored stimuli were not common across modalities. These results indicate that selective attention enhances decodability of shared semantic features across modalities.

The significant intra-modal decoding of latent semantic features in unimodal experiments (**Figure 2**) observed is consistent with previous studies (Pereira et al. 2018; Nishida and Nishimoto 2018; Kivisaari et al. 2019; Fernandino et al. 2022; Frisby et al. 2023), indicating that semantic information can be obtained with sufficient accuracy from brain activity using continuous latent features. The cross-modal decoding further indicates that shared semantic information can be extracted from brain activity, consistent with previous studies reporting shared brain networks for modality-independent semantics processing (Regev et al. 2013; Deniz et al. 2019; Nakai, Yamaguchi, and Nishimoto 2021; Popham et al. 2021; Tang and Huth 2025). The higher decoding accuracies in intra-modal than in cross-modal conditions is likely due to the contribution of stimulus-dependent low-level features to intra-modal decoding. Conversely, cross-modal decoding may more strongly reflect semantic information as it is less stimulus-dependent (Peelen and Downing 2023; Kaplan, Man, and Greening 2015). The strong correlation between decoding accuracies for all combinations of conditions (**Figure 3**) further supports this argument, in that decodable features were common across modalities.

We found higher correlation coefficients of decoding accuracies when the test data matched the modality (**Figures 3B and C**), compared to when the training data matched the modality (**Figures 3D and E**). This asymmetry in the effects of modality matching could be due to the influence on decoding accuracy of low-level stimulus features in the test data. The effect of low-level features in the training data only indirectly affects the decoding accuracy through the weight matrix. In addition, the cross-modal condition where both training and test data modality did not match resulted in the lowest decoding accuracy (**Figure 3F**). In all other cross-modal conditions, either training or test data were of the same modality, so that features related to the physical properties of the stimulus, rather than semantic information, contributed to decoding. Conversely, this all-nonmatch condition could most strongly reflect modality-independent semantic information. In fact, the high correlation of decoding accuracies in this condition supports the existence of modality-independent semantic information in the brain.

In addition, bimodal experiments showed a clear impact of selective attention on decoding accuracy with same modality training and test stimuli (1^st^, 2^nd^, 5^th^, and 6^th^ bar graphs in **Figure 4**). These results are consistent with previous studies reporting that activity in brain semantic circuits is affected by selective attention (Mittag et al. 2013; Wang and He 2014; Moisala et al. 2015; Regev et al. 2019; Nakai, Yamaguchi, and Nishimoto 2021). As those improvements in decoding accuracy were obtained for the same test data (but with different features), these results were not simply produced by stimulus differences. In contrast, the effect of selective attention was absent in the remaining conditions (3^rd^, 4^th^, 7^th^, and 8^th^ bar graphs in **Figure 4**). In these cases, the attended test stimuli were decoded in a cross-modal manner and the ignored stimuli in an intra-modal manner. Considering the higher decoding accuracy in intra-modal than cross-modal conditions in the unimodal experiment (**Figure 2**), it is likely that the effects of modality matching between training and test data and the effects of selective attention canceled each other out.

In bimodal experiments, decodable features were correlated either when training and test modalities were matched or with selective attention to the test data. Correlation was absent when training and test data were unmatched and no attention was paid (**Figure 5D**), suggesting that both modality matching and selective attention determine the decodable features. An observation supported by the unimodal-bimodal correlation results. Decoding accuracies correlated with matched training and test modalities even without selective attention (**Figure 6C, D**) or when selective attention was paid without modality matching (**Figures 6E and F**). However, the correlation was small with unmatched training and test data and no attention was paid (**Figures 6G and H**). These results indicate that both modality matching and selective attention are required to accurately decode semantic information from brain activity.

The cortical visualization of model weights revealed the contributions of shared brain networks for representing semantic information. Activity in perisylvian regions, including the left inferior frontal and superior temporal cortices, has been frequently reported in multimodal experiments (Jobard et al. 2007; Regev et al. 2013; Deniz et al. 2019; Chen et al. 2024), as well as in selective attention experiments on linguistic stimuli (Moisala et al. 2015; Regev et al. 2019; Nakai, Yamaguchi, and Nishimoto 2021). In particular, our previous study demonstrated that these regions store both modality-invariant and attention-selective semantic information (Nakai, Yamaguchi, and Nishimoto 2021); that is, voxels showing high prediction accuracy for both visual and auditory modalities tend to be affected by selective attention. The current findings indicate that these regions also contribute to decoding of semantic information.

One notable difference from the above research, other than the encoding and decoding directions, is the cumulative effect between modality matching and selective attention. In other words, while both properties contributed to decoding accuracy individually, their combination had an even greater impact. Conversely, when the two worked in opposite directions, the decoding performance did not increase as much (e.g., 3^rd^, 4^th^, 7^th^, and 8^th^ bar graphs in **Figure 4**). Note that the current study does not suggest independence of modality-independent and attention-selective semantic processing. How these two processes affect the same perisylvian region in an accumulative manner needs to be investigated in future research.

Several limitations should be noted in the current study. First, we decoded latent semantic features but did not decode specific words such as nouns and verbs. Although a previous study has shown that specific words can be decoded via latent variables (Nishida and Nishimoto 2018), it is unclear whether the effects of modality and selective attention spread to the decoding accuracy of specific words. Second, although we focused only on semantic information using the Wikipedia2Vec model, it is unclear whether the same results can be obtained for decoding of other linguistic information, such as phonology, syntax, and context. Thus, studies aiming to comprehensively investigate the effects of different linguistic information by performing the same analysis using other features are needed.

## Methods

### Participants

In this study, we used the data obtained in (Nakai, Yamaguchi, and Nishimoto 2021). Six healthy participants (referred to as ID01–ID06; ages 22–29; all native Japanese; two females), with normal vision and hearing, participated in the experiments. Participants were all right-handed, as measured using the Edinburgh inventory (Oldfield 1971) (laterality quotients, 62.5–100). This experiment received approval from the Ethics and Safety Committee of the National Institute of Information and Communications Technology, Osaka, Japan. Informed consent was obtained from all participants prior to their participation in the study.

### Stimuli and tasks

We selected 20 narrative stories from the Corpus of Spontaneous Japanese (Maekawa 2003), of which 14 were used during training runs for both Text-only and Speech-only conditions (total, 28 runs). One narrative was used only in the test run of the Text-only condition, another was used only in the test run of the Speech-only condition, two were used only in the test run of the Attend-visual condition (i.e., simultaneously presented in a single run), and another two were used only in the test run of the Attend-audio condition (i.e., simultaneously presented in a single run). All test runs were conducted twice (total, eight runs). We used different narratives during each test run to avoid adaptation to the redundant presentation of the same content.

All narratives were originally recorded in the auditory modality. Sound signals were controlled by their root mean square and only used in the Speech-only, Attend-visual, and Attend-audio conditions. The visual stimuli used for the Text-only, Attend-visual, and Attend-audio conditions were generated by presenting each spoken segment on the center of the screen. The onset of each visual segment matched that of the corresponding segment in the spoken narrative. The average duration of the spoken narratives (mean ± standard deviation [SD]) was 673 ± 70 s.

During the Speech-only condition, participants were asked to fixate on a fixation cross-on the center of the screen and listened to spoken narratives through MRI-compatible ear tips. Meanwhile, during the Text-only condition, participants read the transcribed narratives displayed on the center of the screen, using the rapid serial visual presentation method (Forster 1970). During the Attend-audio condition, participants listened to the spoken narratives through MRI-compatible ear tips and were instructed to ignore the text simultaneously displayed. Participants were asked not to close their eyes and to fixate on the center of the screen. During the Attend-visual condition, participants were instructed to read the transcribed narratives displayed on the center of the screen, while ignoring the simultaneously presented spoken narratives.

At the beginning of each run, 10 s of dummy scans were acquired, during which the fixation cross was displayed; these dummy scans were later omitted from the final analysis to reduce noise. We also obtained 10 s of scans at the end of each run, during which the fixation cross was displayed; these were included in the analyses. In total, 36 fMRI runs were performed for each participant. Among these, 28 runs were used for model training (14 each, under the Speech-only and Text-only conditions) and 8 for model testing (2 each, under the Text-only, Speech-only, Attend-visual, and Attend-audio conditions). For each participant, the experiments were executed over the course of 7 d, with 4–6 runs per day.

Before the fMRI scan, participants were informed that, they would be asked to answer 10 questions relevant to the stimulus they were instructed to concentrate on (the attended stimulus) after each run. However, the actual questionnaire administered after the fMRI scans included 10 questions relevant to both attended and ignored stimuli. The surprise aimed to ensure participants focused on the instructed modality, ignoring distractions.

### MRI data acquisition

This experiment was conducted on a 3.0-T MRI scanner (MAGNETON Prisma; Siemens, Erlangen, Germany) with a 64-channel head coil. We scanned 72 2.0-mm-thick interleaved, axial slices, without a gap, using a T2-weighted, gradient-echo, multiband, echo-planar imaging (MB-EPI) sequence [repetition time (TR) = 1,000 ms, echo time (TE) = 30 ms, flip angle (FA) = 62°, field of view (FOV) = 192 × 192 mm^2^, voxel size = 2 × 2 × 2 mm^3^, multiband factor = 6]. The number of volumes collected varied across runs depending on the stimulus duration, with a mean duration of 693 ± 70 seconds (including 10 seconds of initial dummy scans and 10 seconds of fixation scans at the end of each run). For anatomical reference, high-resolution T1-weighted images of the whole brain were acquired from all participants using a magnetization-prepared rapid acquisition gradient-echo sequence (MPRAGE, TR = 2,530 ms, TE = 3.26 ms, FA = 9°, FOV = 256 × 256 mm^2^, voxel size = 1 × 1 × 1 mm^3^).

### fMRI data preprocessing

Motion correction was performed for each run using the Statistical Parametric Mapping Toolbox (SPM8). All volumes were aligned using the first echo-planar imaging result for each participant. Low-frequency drift was removed using a median filter with a 120-s window. The response for each voxel was then normalized by subtracting the mean response and scaling the response to the unit variance. We used FreeSurfer (Dale, Fischl, and Sereno 1999; Fischl, Sereno, and Dale 1999) to identify cortical surfaces based on anatomical data and to register them against voxels for functional data.

### Semantic features

To quantitatively evaluate the brain representations of the presented semantic information in a data-driven manner, we extracted the semantic features from each narrative stimulus using Wikipedia2Vec (Yamada et al. 2018) (https://wikipedia2vec.github.io/wikipedia2vec/). Wikipedia2Vec has been used to embed words and entities into a distributed representation based on the skip-gram model (Mikolov et al. 2013). The Wikipedia2Vec model is considered an extension of the conventional Word2Vec model, which we previously used (Nishida and Nishimoto 2018). The Word2Vec model is trained solely on contextual words around a target word and has difficulty in dealing with entities (e.g., New York and Julius Caesar). In contrast, the Wikipedia2Vec model is trained on both contextual words and entity link information. All transcribed narrative segments were further segmented into words and morphologically parsed using MeCab (https://taku910.github.io/mecab/). Individual word segments were projected into the 300-dimensional space (i.e., word vectors with 300 elements) and later assigned to the mean time point between onset and offset of target segments, with 40 Hz. The dimension of the word vectors was set to a default value of 300. Time points without any word vector assignments were defined as 0. The resultant concatenated vectors were downsampled to 1 Hz.

### Decoding-model fitting

We separately trained the visual and auditory trained decoding models from Speech-only and Text-only conditions. In the decoding model, the cortical response matrix R [T × 6V] was modeled using concatenating sets of [T × V] matrices with temporal delays of 2–7 s, where T denotes the length of time and V represents the number of voxels. The feature matrix F [T × N] was modeled by multiplying the cortical response matrix R by the weight matrix W [6V × N] (N represents the number of features (=300)):

***F = RW***.

The weight matrix W was estimated using an L2-regularized linear regression with the training dataset. The training dataset comprised 9815 samples under both Speech-only and Text-only conditions. The optimal regularization parameter was assessed in each voxel using 10-fold cross-validation. For each regularization parameter candidate, the decoding accuracy was compared with the data not used to estimate the weights, and the best performing regularization parameter candidate selected. Parameters were searched in the range of 2^0^ to 2^10^. The test dataset used in unimodal analysis comprised 619 Speech-only samples, 617 Text-only samples, and that in bimodal analysis comprised 613 Attend-audio samples, and 623 Attend-visual samples. Differences in sample sizes in the test dataset can be attributed to the various durations of the naturalistic narrative story stimuli. Two repetitions of the test dataset were averaged to increase the signal-to-noise ratio. Pearson’s correlation coefficient between the features of the presented linguistic stimuli and those output by the model was used for decoding accuracy.

### Visualization of model weights

We followed the method of (Nishida and Nishimoto 2018) to visualize the decoding-model weights for each participant. We initially computed average weights across the time lags (six sets), resulting in a V × N matrix, where V represents the number of voxels and N denotes the vector dimensionality (= 300). Voxels exhibiting high decoding weights were considered computational contributors to the decoding process. However, these voxels cannot be simply categorized as indicators of rich perceptual information, as high decoding weights in voxels may result from response covariance across voxels, potentially producing uninformative signals (Haufe et al. 2014). To address this issue, we corrected the decoding-model weights, following the approach proposed by (Haufe et al. 2014): the original weights were left-multiplied by the covariance matrix computed from the voxel response time courses. Then, the absolute values of the corrected weights were taken and the maximum for each voxel selected, leading to an N-dimensional vector. Finally, the vector was z-scored for each participant and projected onto a cortical surface. We considered voxels with higher z-scores more informative.

## Data and code availability

The preprocessed brain data and analysis code used in the current study are available from the corresponding author on reasonable request.

## Acknowledgments

We thank MEXT/JSPS KAKENHI (grant numbers JP24H02172 and JP24H01559 for T.N. and JP18H05522 and JP24H00619 for S.N.), JST ERATO JPMJER1801, AIP JPMJCR24U2 (for S.N.), and the JST FOREST Program (JPMJFR231V for T.N.) for the partial financial support of this study. The funders had no role in the study design, data collection and analysis, decision to publish, or preparation of the manuscript.

## Author contributions

**A.O.**: Conceptualization, Methodology, Formal analysis, Visualization, Writing - Original draft preparation. **H.Y.Q.**: Data collection, Writing - review & editing. **S.N.**: Supervision, Writing - review & editing, Funding acquisition. **T.N.**: Formal analysis, Conceptualization, Methodology, Writing - Original draft preparation, Supervision, Funding acquisition.

## Competing interests

The authors declare no competing interests.

## Supplementary information

**Table S1.** Decoding accuracies and percentage of significant features in unimodal analysis.

|  | ID01 | ID02 | ID03 | ID04 | ID05 | ID06 |
| --- | --- | --- | --- | --- | --- | --- |
| Auditory to auditory | 0.114<br>(68.0%) | 0.138<br>(87.7%) | 0.125<br>(82.0%) | 0.131<br>(82.3%) | 0.146<br>(85.3%) | 0.097<br>(67.0%) |
| Visual to auditory | 0.090<br>(62.3%) | 0.086<br>(54.0%) | 0.089<br>(62.3%) | 0.081<br>(60.7%) | 0.081<br>(54.0%) | 0.074<br>(47.3%) |
| Visual to visual | 0.203<br>(90.7%) | 0.204<br>(91.7%) | 0.159<br>(84.7%) | 0.119<br>(74.0%) | 0.215<br>(93.3%) | 0.126<br>(78.7%) |
| Auditory to visual | 0.125<br>(76.0%) | 0.126<br>(75.7%) | 0.088<br>(57.0%) | 0.093<br>(63.3%) | 0.135<br>(82.0%) | 0.094<br>(64.3%) |

Decoding accuracy evaluated using Pearson's correlation coefficient ( $r$ ). Percentage of significant features in parentheses.

**Table S2.** Comparison of decoding accuracies in unimodal experiments.

|  | ID01 | ID02 | ID03 | ID04 | ID05 | ID06 |
| --- | --- | --- | --- | --- | --- | --- |
| Visual to visual against auditory to auditory | 0.695 | 0.608 | 0.582 | 0.612 | 0.777 | 0.581 |
| Visual to visual against auditory to visual | 0.773 | 0.767 | 0.651 | 0.665 | 0.820 | 0.719 |
| Auditory to auditory against visual to auditory | 0.468 | 0.547 | 0.696 | 0.567 | 0.656 | 0.638 |
| Visual to visual against visual to auditory | 0.363 | 0.327 | 0.435 | 0.382 | 0.628 | 0.496 |
| Auditory to auditory against auditory to visual | 0.600 | 0.582 | 0.454 | 0.605 | 0.657 | 0.540 |
| Visual to auditory against auditory to visual | 0.349 | 0.447 | 0.435 | 0.449 | 0.564 | 0.443 |

Relation between decoding accuracies evaluated using Spearman's correlation coefficient ( $\rho$ ).

**Table S3.** Decoding accuracies and percentage of significant features in unimodal analysis.

|  | ID01 | ID02 | ID03 | ID04 | ID05 | ID06 |
| --- | --- | --- | --- | --- | --- | --- |
| Auditory-to-auditory (attend) | 0.103<br>(61.3%) | 0.170<br>(86.0%) | 0.137<br>(76.3%) | 0.147<br>(78.0%) | 0.159<br>(88.7%) | 0.115<br>(70.0%) |
| Auditory to visual (ignored) | -0.002<br>(1.7%) | 0.011<br>(0.7%) | -0.002<br>(1.0%) | 0.016<br>(1.3%) | 0.006<br>(0.7%) | 0.025<br>(5.7%) |
| Visual to auditory (attend) | 0.069<br>(46.0%) | 0.090<br>(65.0%) | 0.103<br>(71.0%) | 0.080<br>(57.7%) | 0.104<br>(73.0%) | 0.102<br>(64.7%) |
| Visual to visual (ignored) | 0.082<br>(55.7%) | 0.095<br>(62.0%) | 0.057<br>(39.3%) | 0.053<br>(34.0%) | 0.089<br>(58.7%) | 0.041<br>(20.7%) |
| Visual to visual (attend) | 0.158<br>(82.3%) | 0.197<br>(92.7%) | 0.109<br>(76.3%) | 0.144<br>(84.7%) | 0.225<br>(94.0%) | 0.118<br>(77.0%) |
| Visual to auditory (ignored) | 0.043<br>(12.3%) | 0.038<br>(8.3%) | 0.061<br>(44.3%) | 0.029<br>(7.3%) | 0.028<br>(5.3%) | 0.048<br>(26.7%) |
| Auditory to visual (attend) | 0.047<br>(36.0%) | 0.080<br>(59.3%) | 0.037<br>(7.7%) | 0.067<br>(49.3%) | 0.082<br>(63.7%) | 0.079<br>(58.0%) |
| Auditory to auditory (ignored) | 0.073<br>(51.0%) | 0.107<br>(66.7%) | 0.099<br>(61.3%) | 0.118<br>(69.7%) | 0.1012<br>(63.7%) | 0.087<br>(60.7%) |

Decoding accuracy evaluated using Pearson's correlation coefficient ( $r$ ). Percentage of significant features in parentheses.

**Table S4.** Comparison of decoding accuracies in bimodal experiments.

|  | ID01 | ID02 | ID03 | ID04 | ID05 | ID06 |
| --- | --- | --- | --- | --- | --- | --- |
| Visual to visual (attend)<br>against auditory to auditory (attend) | 0.744 | 0.759 | 0.586 | 0.642 | 0.698 | 0.540 |
| Visual to visual (ignored)<br>against auditory to auditory (ignored) | 0.697 | 0.702 | 0.488 | 0.451 | 0.586 | 0.445 |
| Visual to auditory (attend)<br>against auditory to visual (attend) | 0.414 | 0.425 | 0.280 | 0.360 | 0.357 | 0.442 |
| Visual to auditory (ignored)<br>against auditory to visual (ignored) | 0.161 | -0.059 | 0.019 | 0.160 | 0.037 | 0.304 |

Relation between decoding accuracies evaluated using Spearman's correlation coefficient ( $\rho$ ).

**Table S5.**
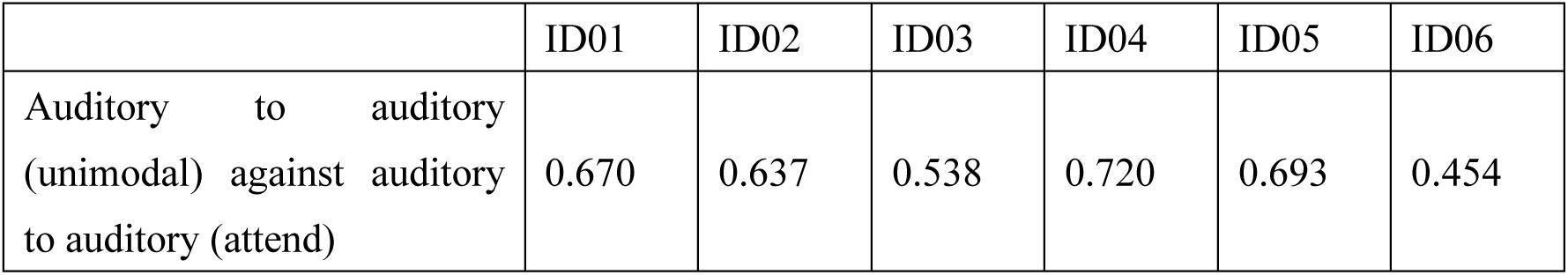
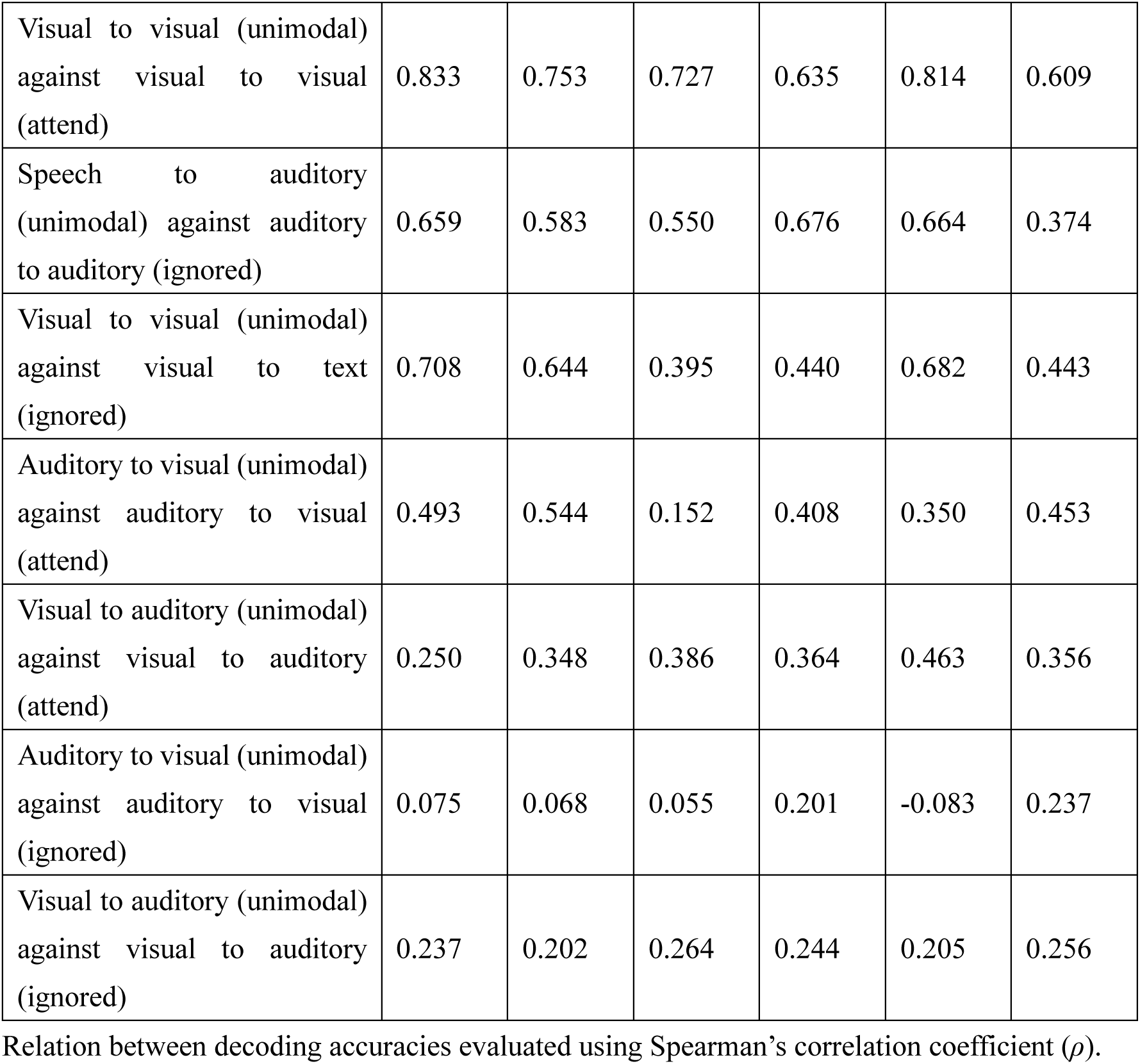
Comparison of decoding accuracies between unimodal and bimodal experiments.

|  | ID01 | ID02 | ID03 | ID04 | ID05 | ID06 |
| --- | --- | --- | --- | --- | --- | --- |
| Auditory to auditory (unimodal) against auditory to auditory (attend) | 0.670 | 0.637 | 0.538 | 0.720 | 0.693 | 0.454 |
| Visual to visual (unimodal)<br>against visual to visual<br>(attend) | 0.833 | 0.753 | 0.727 | 0.635 | 0.814 | 0.609 |
| Speech to auditory<br>(unimodal) against auditory<br>to auditory (ignored) | 0.659 | 0.583 | 0.550 | 0.676 | 0.664 | 0.374 |
| Visual to visual (unimodal)<br>against visual to text<br>(ignored) | 0.708 | 0.644 | 0.395 | 0.440 | 0.682 | 0.443 |
| Auditory to visual (unimodal)<br>against auditory to visual<br>(attend) | 0.493 | 0.544 | 0.152 | 0.408 | 0.350 | 0.453 |
| Visual to auditory (unimodal)<br>against visual to auditory<br>(attend) | 0.250 | 0.348 | 0.386 | 0.364 | 0.463 | 0.356 |
| Auditory to visual (unimodal)<br>against auditory to visual<br>(ignored) | 0.075 | 0.068 | 0.055 | 0.201 | -0.083 | 0.237 |
| Visual to auditory (unimodal)<br>against visual to auditory<br>(ignored) | 0.237 | 0.202 | 0.264 | 0.244 | 0.205 | 0.256 |

Relation between decoding accuracies evaluated using Spearman's correlation coefficient ( $\rho$ ).

**Figure S1.**
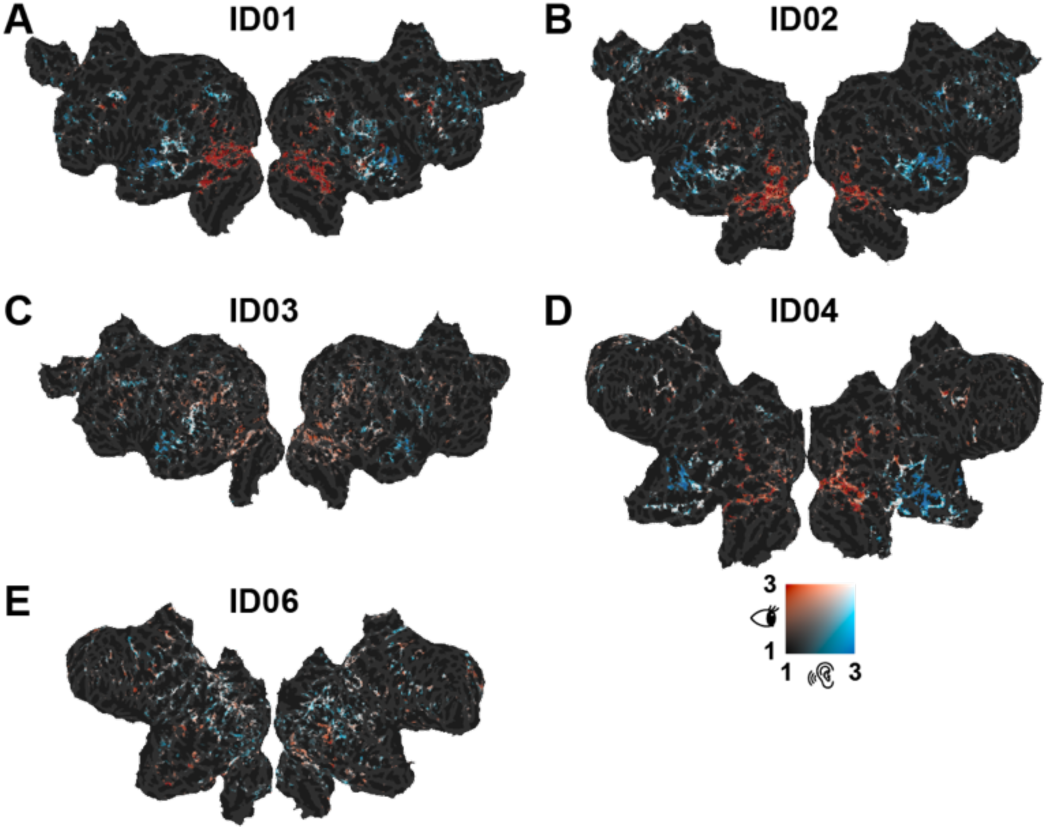
Contributing voxels for other participants. Maximum absolute model weights projected onto the cortical surface of participants ID01, ID02, ID03, ID04, and ID06. Red voxels correspond to the visual model and blue voxels to the auditory model.

**Figure S2.**
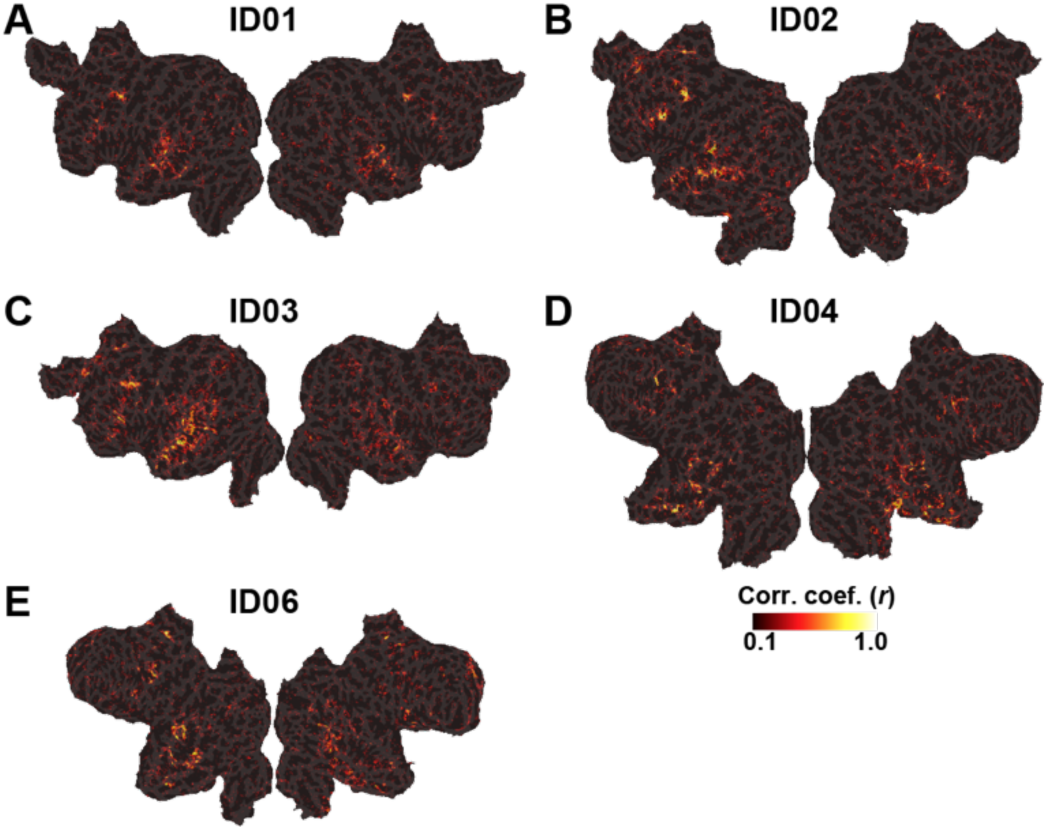
Shared voxels across modalities for other participants. Correlations between the visual and auditory model weights projected onto the cortical surface of participants ID01, ID02, ID03, ID04, and ID06.

## Notes

### Competing Interest Statement

The authors have declared no competing interest.

